# Organ-specific Sympathetic Innervation Defines Visceral Functions

**DOI:** 10.1101/2024.09.19.613934

**Authors:** Tongtong Wang, Bochuan Teng, Dickson R. Yao, Wei Gao, Yuki Oka

## Abstract

The autonomic nervous system orchestrates the brain and body functions through the sympathetic and parasympathetic pathways. However, our understanding of the autonomic system, especially the sympathetic system, at the cellular and molecular levels is severely limited. Here, we show unique topological representations of individual visceral organs in the major abdominal sympathetic ganglion complex. Using multi-modal transcriptomic analyses, we identified distinct sympathetic populations that are both molecularly and spatially separable in the celiac-superior mesenteric ganglia (CG-SMG). Notably, individual CG-SMG populations exhibit selective and mutually exclusive axonal projections to visceral organs, targeting either the gastrointestinal (GI) tract or secretory areas including the pancreas and bile tract. This combinatorial innervation pattern suggests functional segregation between different CG-SMG populations. Indeed, our neural perturbation experiments demonstrated that one class of neurons selectively regulates GI food transit. Another class of neurons controls digestion and glucagon secretion independent of gut motility. These results reveal the molecularly diverse sympathetic system and suggest modular regulations of visceral organ functions through distinct sympathetic populations.

## Introduction

The brain-body bidirectional communication via sensory and autonomic pathways is crucial in regulating various physiological functions ranging from nutrient digestion^1^, and cardiovascular functions^2,3^, to thermoregulation^4,5^. Recent studies demonstrated the functional diversity of sensory vagal and spinal neurons transmitting various types of body-to-brain signals^6–11^. The descending autonomic responses are controlled by two primary nervous systems: the sympathetic and parasympathetic systems, which use norepinephrine and acetylcholine as neurotransmitters^12^.

The sympathetic nervous system in vertebrate animals is activated under stress and mediates adrenergic “fight or flight” response by increasing heart rate and blood glucose level while decreasing digestion and body fluid secretion^13–16^. The parasympathetic system counteracts the sympathetic signals^12^. A few studies recently identified genetically distinct parasympathetic pathways to cardiovascular^2^ and gastrointestinal^17^ areas. However, the cellular and functional organization of the sympathetic nervous system is still poorly understood. While the anatomy of sympathetic projections has been well-described^18–21^, how sympathetic neurons regulate visceral functions remains unknown. The lack of basic understanding poses a challenge in developing therapeutic interventions for autonomic disorders such as bowel syndrome and gastroparesis^22^. The current treatments still rely on non-specific modulation of neurotransmitters identified over a century ago.

In this study, we applied transcriptomics, organ mapping, and cell-type-specific functional perturbation to the major visceral sympathetic ganglia, CG-SMG. Our results demonstrate that molecularly distinct sympathetic neuron types innervate unique combinations of visceral organs, working independently to regulate food transit and secretory functions.

## Results

Peripheral organs receive sympathetic inputs from pre- and para-vertebral ganglia organized in a topological manner known as the sympathetic chain^23^. The CG-SMG complex consists of three ganglia (bilateral CG and SMG) that send sympathetic signals to visceral organs^24^. To gain insights into sympathetic regulatory mechanisms, we determined the topological representation of visceral organs in CG-SMG (Fig. 1). We performed retrograde tracing using wheat germ agglutinin (WGA) from eight visceral organs: spleen, hepatic portal area (HPA), pancreas, stomach, duodenum, jejunum, ileum, and colon. We developed a computational pipeline to build a standardized CG-SMG atlas for precise ganglia-organ mapping across animals (Fig. 1a, Extended Data Fig. 1a,b). The reference anatomy was generated from 20 ganglia through unsupervised image alignment processing. We then mapped auto-detected WGA signals onto the reference image (Extended Data Fig. 1c). This analysis revealed unique topographic representations of visceral organs on CG-SMG (Fig. 1b, Extended Data Fig. 2a-c). Neurons that project to the spleen, HPA, pancreas, stomach, and duodenum were predominantly located at lateral CG-SMG. Distal small intestine and large intestine were mainly mapped to medial areas. We next asked whether different organs receive inputs from the same or different sets of CG-SMG neurons. Consistent with a recent study on the GI tract^25^, visceral organs received innervation from non-overlapping neurons. We performed anatomical tracing from paired organ sites by dual-color WGA labeling. While injection of both WGAs in the same organ produced mostly overlapping neural labeling, we found minimal overlap of WGA signals derived from two different organs (Fig. 1c, Extended Data Fig. 2d). These results revealed that adjacent organs tend to be represented in proximal areas of CG-SMG while receiving inputs from different sets of neurons.

**Fig. 1:**
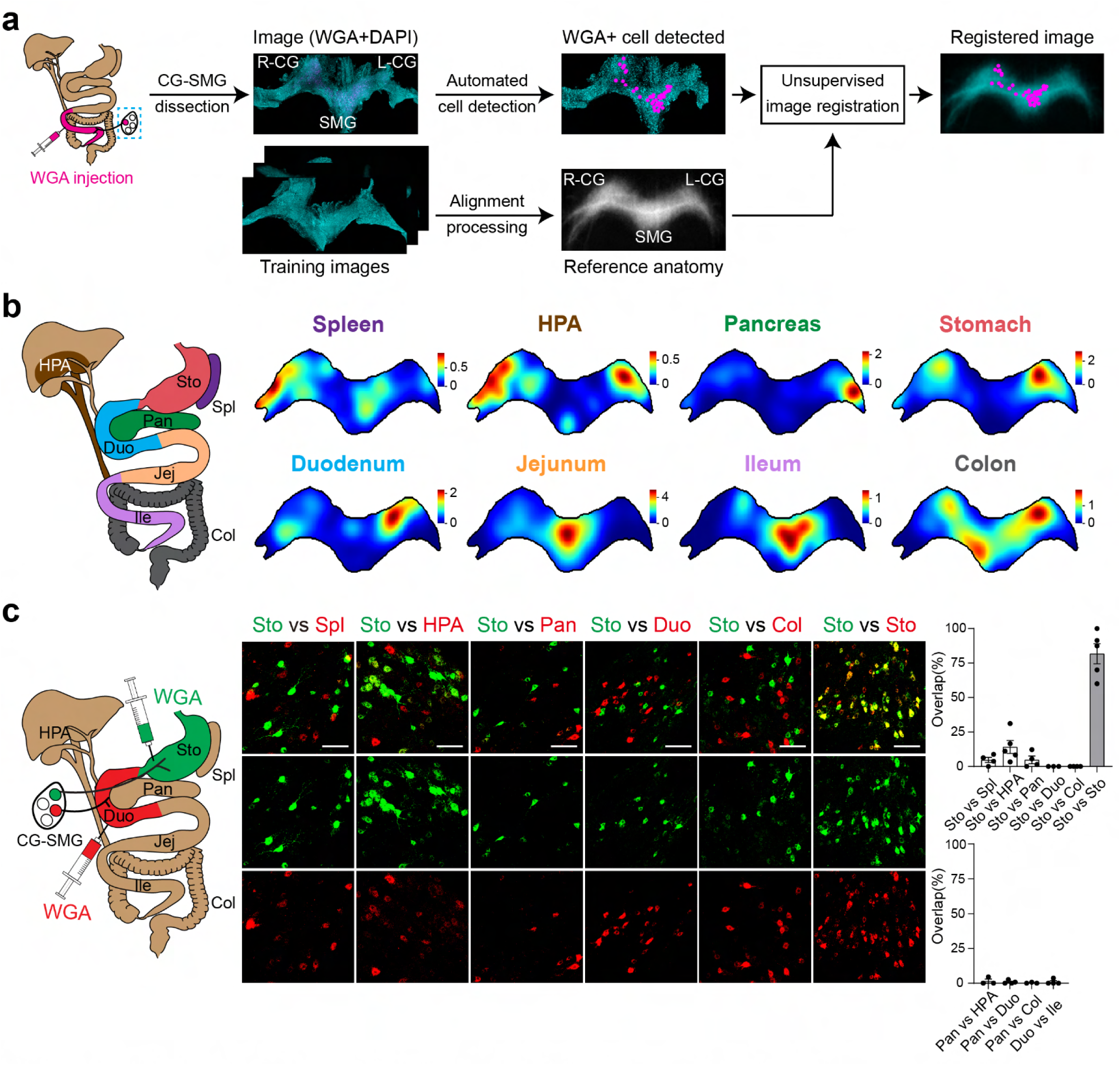
Topological representation of visceral organs in CG-SMG sympathetic complex. **a,** A computational pipeline for organ topographical mapping. WGA was injected in a given visceral organ for retrogradely labeling CG-SMG neurons. WGA-positive cells (magenta) were auto-detected on CG-SMG using DAPI (blue) as a background shape. For aligning ganglia across animals, 20 training images were used to build a standardized reference image through an unsupervised alignment process. Organ maps (WGA-positive neurons) from each animal were registered to the reference atlas. **b**, Gaussian heatmaps of WGA-positive cells traced from individual visceral organs. Eight organ sites were targeted: the spleen (Spl), hepatic portal area (HPA), pancreas (Pan), stomach (Sto), duodenum (Duo), jejunum (Jej), ileum (Ile), and colon (Col). For each organ site, the average of heatmaps from three mice is presented. **c,** WGA dual-color tracing experiments of visceral organ pairs. Left: a diagram showing WGA 488 (green) injection to the stomach and WGA 647 (red) injection to the duodenum. Middle: representative CG-SMG images with two-color WGA injections to organs as indicated. Minimal overlap was detected between paired organs. Right: quantification of the overlapped neurons as a percentage of total WGA-positive neurons. n = 4, 5, 4, 3, 4, 5 mice for Sto vs Spl, Sto vs HPA, Sto vs Pan, Sto vs Duo, Sto vs Col, and Sto vs Sto. n = 3, 4, 3, 4 mice for Pan vs HPA, Pan vs Duo, Pan vs Col, and Duo vs Ile. Scale bar, 100 μm. Data are presented as mean ± s.e.m.

We next used transcriptomic approaches to examine the cellular diversity of CG-SMG neurons. Toward this goal, we performed single-cell RNA sequencing (scRNA-seq) from 25k CG-SMG cells. Based on uniquely expressed genes in each cell cluster, we selected 371 genes for spatial transcriptomics experiments (Fig. 2a, Extended Data Fig. 3a). Freshly frozen CG-SMG samples were subjected to SeqFISH^26^ to visualize spatial gene expression map. We identified tyrosine hydroxylase (TH)-positive neurons and other non-neuronal cell types (Extended Data Fig. 3b)^27–29^. Further analyses focusing on TH-positive CG-SMG (CG-SMG^TH^) neurons revealed two molecularly distinct neuron classes expressing relaxin receptor type 1 (CG-SMG^RXFP1^) or a homeobox gene, SHOX2 (CG-SMG^SHOX2^, Fig. 2b). Other neuropeptides described previously (e.g., NPY) were broadly expressed by CG-SMG neurons (Extended Data Fig. 3c)^30–33^. We also found that CG-SMG^SHOX2^ neurons comprise at least two subclasses labeled by the expression of *BMP3* and *DSP* (Fig. 2c). However, compared to sensory ganglia and parasympathetic nuclei^2,17,34–36^, CG-SMG neurons exhibited less diversity, presumably reflecting the simplicity of sympathetic regulation of visceral organs.

**Fig. 2:**
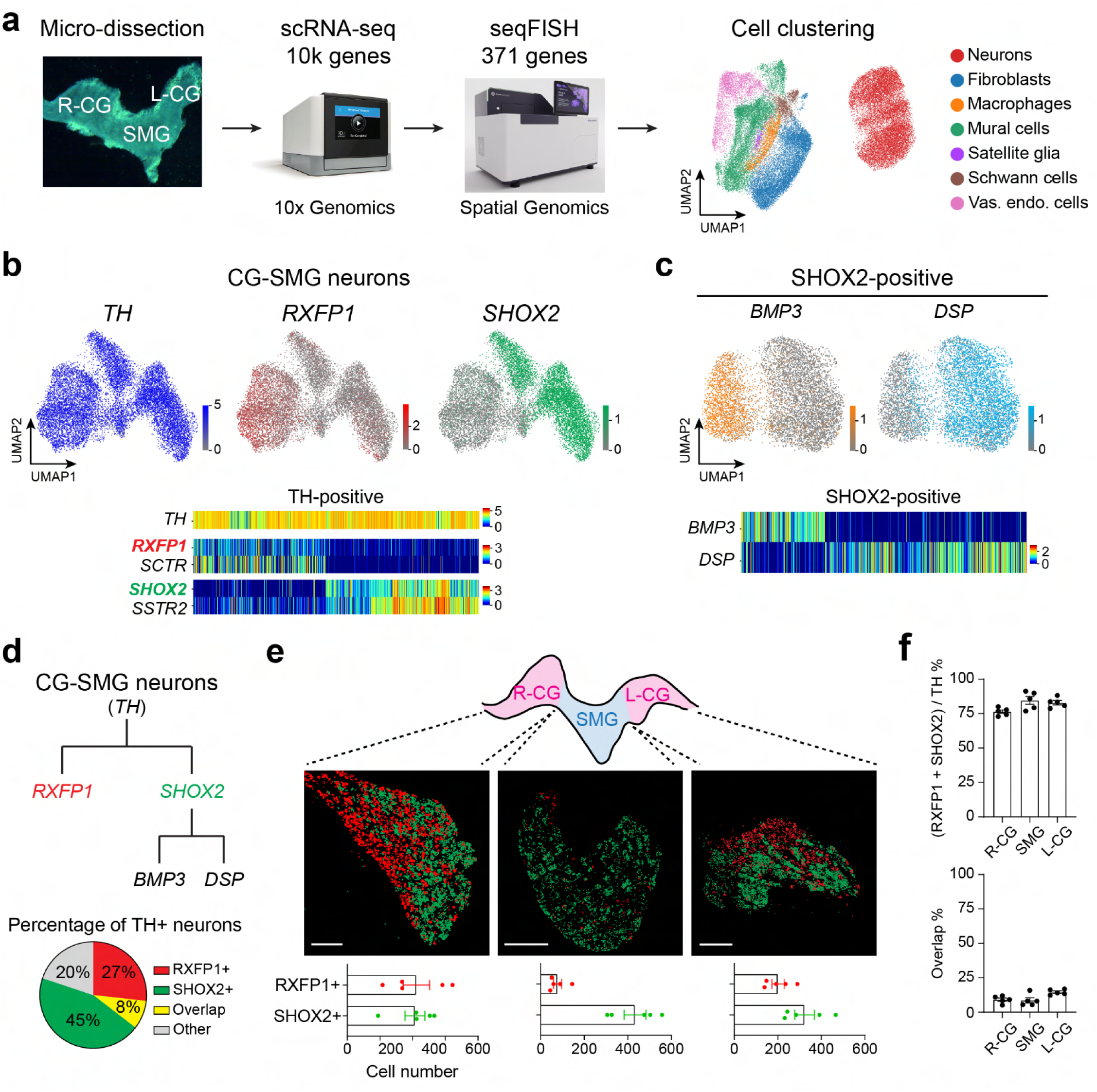
Multi-transcriptomic analyses reveal molecularly and spatially distinct CG-SMG neural populations. **a,** Experimental design for identifying transcriptomic cell types and for visualizing spatial localization of cells in the CG-SMG complex. By analyzing >10k genes in scRNA-seq, 371 cell-type marker genes were used for seqFISH analysis. UMAP embedding of major cell types identified from seqFISH experiments (n = 25,932 cells from 5 sections of 3 mice per Right (R)-CG, SMG, Left (L)-CG). Vas. endo. cells, vascular endothelial cells. **b,** SeqFISH analysis of CG-SMG neurons. Top: UMAP plotting for log-normalized *TH* (blue), *RXFP1* (red) and *SHOX2* (green) expression. Bottom: heatmaps of neuronal marker gene expression (n = 13,105 cells). **c,** UMAP (top) and heatmap (bottom) plotting for subtypes of CG-SMG^SHOX2^ neurons (n = 7,152 cells). **d,** Top: a summary diagram of CG-SMG neuronal types. Bottom: a pie chart displaying the percentage of neuron types. **e,** Spatial localization of CG-SMG^RXFP1^ and CG-SMG^SHOX2^ neurons. Representative seqFISH images for the expression of *RXFP1* (red) and *SHOX2* (green) in CG-SMG areas. Scale bar, 200 μm. Shown are quantification of RXFP1- and SHOX2-positive neurons (n = 5 sections of 3 mice per R-CG, SMG, L-CG). **f,** Quantification of RXFP1- and SHOX2-positive neurons in each area of CG-SMG (top) and overlapping percentage (bottom, n = 5 sections of 3 mice per R-CG, SMG, L-CG). Data are presented as mean ± s.e.m.

CG-SMG^RXFP1^ and CG-SMG^SHOX2^ neurons account for −80 % of the entire CG-SMG^TH^ neurons with minimal overlap (Fig. 2d). These two complementary populations showed unique topological distribution in CG-SMG: CG-SMG^SHOX2^ neurons were distributed across both CG and SMG whereas the vast majority of CG-SMG^RXFP1^ neurons were located only in CG (Fig. 2e,f). Moreover, different subclasses of CG-SMG^SHOX2^ neurons were intermingled in both CG and SMG (Extended Data Fig. 3d,e).

Given distinct CG-SMG neuron types, we explored the neuron-to-organ relationship. To label individual CG-SMG populations, we used RXFP1-Cre and SHOX2-Cre^37^ animals. In both transgenic lines, we confirmed that Cre was faithfully expressed in endogenous CG-SMG^RXFP1^ and CG-SMG^SHOX2^ neurons (Extended Data Fig. 4). For virus tracing, each transgenic animal was injected with AAV-FLEX-tdTomato in the whole CG-SMG complex (Fig. 3a). After 3 weeks of recovery, each organ was dissected, cleared in ScaleS solution^38^, and fluorescent projection was visualized. In all animals, we validated robust fluorescent expression in CG-SMG (Extended Data Fig. 5a,b). TH-Cre animals were used to target CG-SMG neurons ubiquitously.

**Fig. 3:**
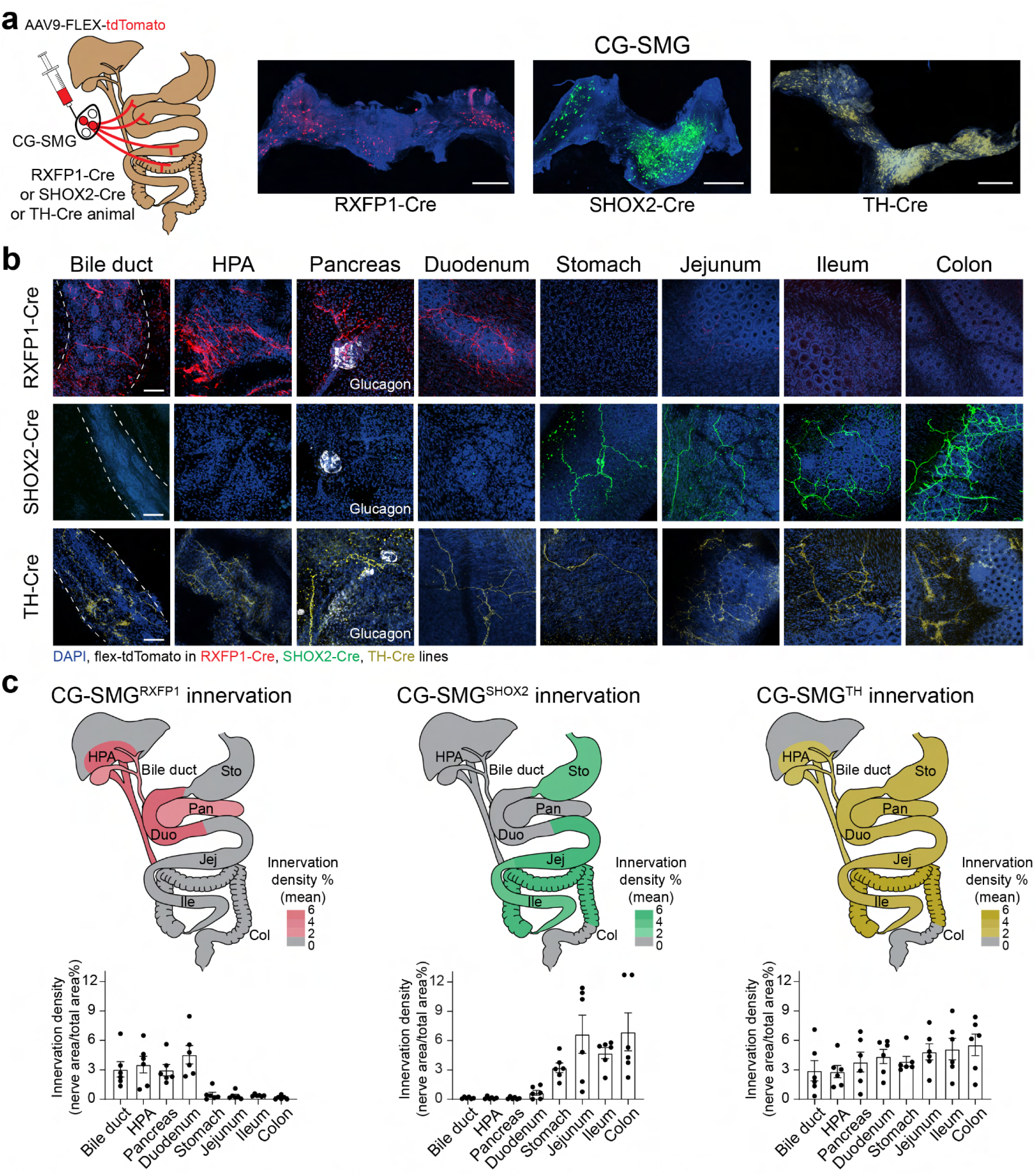
Complementary organ projections by CG-SMG^RXFP1^ and CG-SMG^SHOX2^ neurons. **a,** CG-SMG neuron anterograde tracing. Cell bodies and nerve terminals were visualized by injecting AAV-FLEX-tdTomato into CG-SMG of RXFP1-Cre, SHOX2-Cre and TH-Cre animals. Representative images of tdTomato expression after AAV infection are shown with TH immunostaining (blue). Scale bar, 500 μm. **b,** Visualization of CG-SMG axons in visceral organs. Representative images for whole-mount organ innervation of CG-SMG^RXFP1^, CG-SMG^SHOX2^, CG-SMG^TH^ neurons as indicated (n = 3). Top and middle: RXFP1+ and SHOX2+ neurons target complementary visceral organs. Bottom: TH+ neurons project to all visceral organs as a positive control. The pancreatic islet is identified by the indicated glucagon antibody staining (white) and nuclei are visualized by DAPI staining (blue). Scale bar, 100 μm. **c,** Summary maps of anatomical innervation patterns by CG-SMG neurons with color-coded average nerve density. The nerve density quantification is shown at the bottom (n = 6 images from 3 mice per organ). HPA, hepatic portal area. Pan, pancreas. Duo, duodenum. Sto, stomach. Jej, jejunum. Ile, ileum. Col, colon. Data are presented as mean ± s.e.m.

Surprisingly, our projection analyses identified that each CG-SMG population innervates unique combinations of visceral organs in a mutually exclusive fashion (Fig. 3b,c and Extended Data Fig. 5c,d). CG-SMG^RXFP1^ neurons sent projections to the upper digestive and secretory organs, including the bile duct, HPA, pancreas, and duodenum. The stomach was devoid of projections from CG-SMG^RXFP1^ neurons. Conversely, CG-SMG^SHOX2^ neurons projected to the stomach, and lower intestinal areas (jejunum, ileum, and colon). All visceral organs received strong innervations from CG-SMG^TH^ neurons. These combinatorial organ projection patterns from individual CG-SMG neuron types led us to speculate that each population may mediate specific visceral functions.

Recent transcriptomic studies on peripheral ganglia have provided insights into the cellular composition of the autonomic system^27,28^. Nevertheless, the specific functions of each neuronal classes remains unclear. To fill this gap, we applied functional perturbation tools to CG-SMG neuron types *in vivo*. We first electrophysiologically characterized action potentials induced by optogenetic and chemogenetic stimulation. TH-Cre;Ai32 double transgenic animals were used to express channelrhodopsin (ChR2) in CG-SMG^TH^ neurons (Fig. 4a). CG-SMG neurons showed action potentials in response to light pulses, and increasing light frequency caused higher firing rate which plateaued around 10 Hz (Fig. 4b). For chemogenetic stimulation, CG-SMG of TH-Cre animals were injected with AAV-FLEX-hM3Dq. Clozapine *N*-oxide (CNO) application to DREADD-expressing CG-SMG neurons induced persistent firing (Fig. 4c). These results validated functional perturbation tools in CG-SMG neurons.

**Fig. 4:**
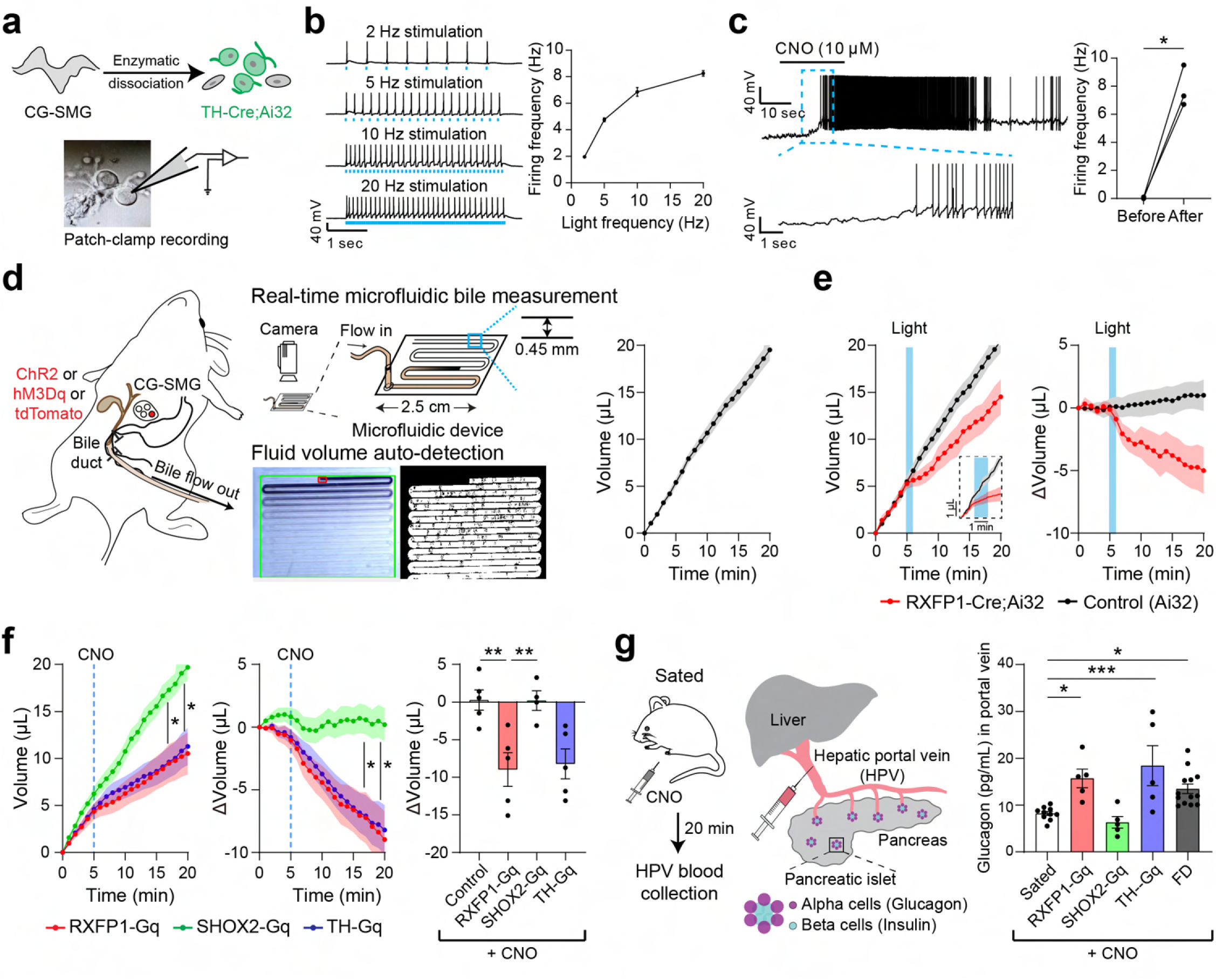
Activation of CG-SMG^RXFP1^ neurons regulates secretory functions. **a,** Patch clamp electrophysiological recording from dissociated ChR2-expressing CG-SMG neurons. **b,** Membrane potential responses of a ChR2-transfected CG-SMG neuron to repetitive trains of 10 ms-long light pulses as indicated. Quantification of firing frequency at different light frequencies is shown (n = 5 neurons from 3 mice). **c,** Validation of chemogenetic manipulation of CG-SMG neurons. Left: action potentials upon CNO application (10 µM). Right: quantification of firing rate before and after CNO application (n = 3 neurons from 2 mice). **d,** A microfluidic device for automated measurement of bile secretion *in vivo*. Left and middle: schematic for measuring bile secretion volume. Bile flow was connected to the custom microfluidic chamber and was recorded by a camera. Right: Automated fluid detection of bile volume across time is plotted (n = 9 mice). **e,** Optogenetic stimulation of CG-SMG^RXFP1^ neurons inhibited bile secretion in a time-lock manner. Total bile secretion (left) and volume change (right) over time (n = 4 RXFP1-Cre;Ai32 mice, n = 6 cagemate control mice). Light pulses of 2 ms at 20 Hz were applied at CG-SMG for 1 min as indicated by the blue shade. A zoomed-in view shows a 3-min peri-stimulation window with 30 Hz sampling rate. **f,** Effects of chemogenetic activation of CG-SMG neurons on bile secretion. Left: Chemogenetic activation of CG-SMG^RXFP1^ (red) and CG-SMG^TH^ (blue) neurons significantly suppressed bile secretion, but not CG-SMG^SHOX2^ (green) neurons. Middle: Bile volume change upon chemogenetic stimulation. Right: Quantification of volume change after chemogenetic stimulation. RXFP1-Cre animals injected with tdTomato were used as a control group. Data were quantified from 5 mice for RXFP1-tdT, 5 mice for RXFP1-Gq, 4 mice for SHOX2-Gq, and 5 mice for TH-Gq. **g,** Effects of chemogenetic activation of CG-SMG neurons on glucagon release. Left and Middle: Experimental protocol of measuring blood glucagon level. Right: Activation of CG-SMG^RXFP1^ (red), but not CG-SMG^SHOX2^ neurons (green) increased glucagon release under sated conditions. n = 10, 5, 5, 5, 13 mice for wild-type sated, RXFP1-Gq, SHOX2-Gq, TH-Gq, wild-type food-deprived (FD) conditions. *P < 0.05, **P < 0.01, ***P < 0.001 by two-tailed paired t-test, one-way or two-way ANOVA with Dunnett’s multiple comparisons test. Data are presented as mean ± s.e.m.

The visceral sympathetic system regulates various physiological functions, including gut motility and digestion^39^. We tested how individual functions are controlled by CG-SMG^RXFP1^ and CG-SMG^SHOX2^ populations. For digestive functions, we focused on bile secretion, a key component of fat digestion in the intestine that receives sympathetic modulation^40^. Due to the extremely low volume of bile secretion, we developed an automated *in vivo* bile measurement system by combining a microfluidic device and a computer-vision-based system that allows sub-second quantification of bile (Fig. 4d, Extended Data Fig. 6a, b). Under undisturbed conditions, bile secretion was about 1 μl per min (Extended Data Fig. 6c). We explored how secretion speed changes upon manipulation of distinct CG-SMG neurons. Each CG-SMG population was activated either by topical application of CNO or photostimulation (488 nm). As suspected from the projection pattern, optogenetic stimulation of CG-SMG^RXFP1^ neurons significantly slowed down bile secretion compared to a control group (Fig. 4e, Extended Data Fig. 6d). Similar results were obtained with chemogenetic activation (Fig. 4f). Interestingly, CG-SMG^RXFP1^ neurons densely innervate the bile entry site (Extended Data Fig. 5c) suggesting that regulation of bile is mainly at the intestinal entry site (Extended Data Fig. 6e-h).

Our anatomical study showed that the CG-SMG^RXFP1^ neurons also project to other secretory organs, in particular, the pancreatic islet (Extended Data Fig. 5c). Since sympathetic activation leads to glucagon secretion and elevated glucose levels under stress conditions^41,42^, we tested whether the CG-SMG^RXFP1^ population concurrently regulate both bile and glucagon secretion. Under food-deprived conditions, glucagon levels were significantly increased in portal vein blood (Fig. 4g, Extended Data Fig. 6i). Chemogenetic activation of CG-SMG^RXFP1^ neurons also significantly increased glucagon secretion comparable to CG-SMG^TH^ neuron stimulation (Fig. 4g, Extended Data Fig. 6j). By sharp contrast, the same functional manipulation of CG-SMG^SHOX2^ population had no effects on bile or glucagon secretion. These results demonstrate that CG-SMG^RXFP1^ neurons innervate combinatorial upper visceral organs, including the bile duct, duodenum, pancreas, and liver, to control digestion- and secretion-related functions.

We next explored the functions of the CG-SMG^SHOX2^ population that projects complementary organs. These neurons innervate both the muscle layer and myenteric plexus of the stomach and intestine (Fig. 3c, Extended Data Fig. 5d). Since GI organs are heavily involved in gut transit and nutrient absorption^43^, we suspected that the CG-SMG^SHOX2^ population may regulate the GI functions independent of digestion. To test this idea, we quantified food transit by subjecting animals to food deprivation, allowing them brief access to colored chow (10 min), and examining the GI transit (Fig. 5a, Extended Data Fig. 7a). Interestingly, stimulation of CG-SMG^SHOX2^ or CG-SMG^TH^ neurons drastically reduced food transit (Fig. 5b). Conversely, we observed negligible effects when CG-SMG^RXFP1^ neurons were activated (Fig. 5c). We note that the observed difference in transit was not due to reduced food intake (Extended Data Fig. 7b). These results show that CG-SMG^SHOX2^, but not CG-SMG^RXFP1^ neurons control food transit after ingestion. We further tested fecal output in awake-behaving animals as a proxy for lower gut motility^44^. Under sated conditions, CG-SMG^SHOX2^ or CG-SMG^RXFP1^ neurons were activated by CNO while observing the number of stools produced during the next 3-hour period (Fig. 5d). We found that stool expulsion was essentially abolished when CG-SMG^SHOX2^ neurons were activated. By contrast, stimulation of CG-SMG^RXFP1^ neurons did not have any effects (Fig. 5e,f, Extended Data Fig. 7c,d). Collectively, our cell-type-specific manipulation shows that two CG-SMG neural populations are molecularly, spatially, and functionally distinct. This study suggests that sympathetic neurons comprise diverse cell types, each serving as a functional module that controls functionally related visceral organs (Fig. 5g).

**Fig. 5:**
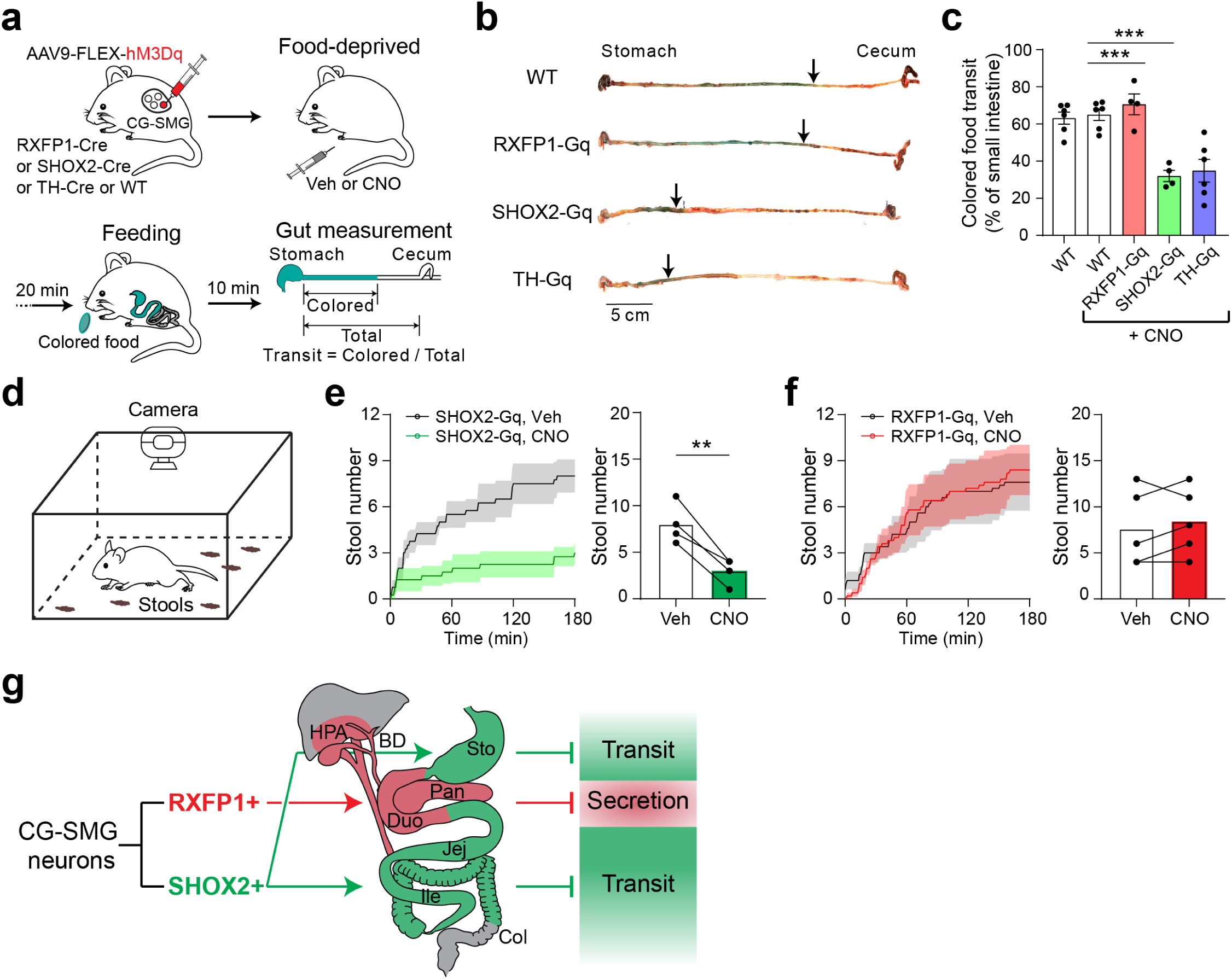
Activation of CG-SMG^SHOX2^ neurons inhibits gut transit. **a,** An experimental design for food transit measurement with chemogenetic activation. **b,** Representative images of the small intestine after activating individual CG-SMG neuron populations. Arrows indicate where colored food (blue) travels after 10 min of feeding. **c,** Quantified data showing that food transit was significantly suppressed by the activation of CG-SMG^SHOX2^ or CG-SMG^TH^ neurons, but not by CG-SMG^RXFP1^ neurons. Wild-type sham-surgery animals were examined after either vehicle (WT-PBS) or CNO (WT-CNO) administration. n = 6 mice for WT-PBS, 6 mice for WT-CNO, 4 mice for RXFP1-Gq, 4 mice for SHOX2-Gq, 6 mice for TH-Gq groups. **d,** Schematic for stool number measurement. Upon CNO application, sated animals were placed in an open field chamber for 3-hour video recording. **e,** Chemogenetic stimulation of CG-SMG^SHOX2^ neurons suppressed spontaneous stool expulsion. The total number of stools was plotted and quantified (n = 4 mice). **f,** Conversely, activation of CG-SMG^RXFP1^ neurons had no effects on stool defecation (n = 5 mice). **g,** Modular sympathetic regulation of visceral organs. **P < 0.01, ***P < 0.001 by two-tailed paired *t*-test or one-way ANOVA with Dunnett’s multiple comparisons test. All data are shown as mean ± s.e.m.

## Discussion

### Cellular diversity of CG-SMG neurons

Recent studies highlighted diverse neural populations in ascending and descending nervous systems. The vagal and spinal sensory neurons comprise molecularly distinct populations with various sensory functions. Similarly, the sympathetic nervous system also shows cellular and molecular diversity. *RXFP1* and *SHOX2* are expressed by two types of neurons spatially segregated in CG-SMG. Furthermore, GI-innervating CG-SMG^SHOX2^ neurons contain at least two populations (Fig. 2d). These results are in accordance with recent studies in other sympathetic chain ganglia^27,28^. However, our transcriptomic analyses suggested less diversity in CG-SMG neurons compared to visceral sensory systems. It is feasible that a less diverse cellular composition may reflect the relatively narrow functionality of the sympathetic system. Alternatively, future higher-resolution transcriptomic analyses may reveal more granular cell classes with distinct functionality.

### Modular functional regulation by the sympathetic system

CG-SMG^RXFP1^ and CG-SMG^SHOX2^ neurons showed complementary projection patterns to visceral organs. The former projects to secretory sites (bile duct, duodenum, pancreas, and the liver), whereas the latter sends innervations to the visceral GI tract. Interestingly, specific visceral functions such as digestion and food transit appear to be controlled by distinct CG-SMG populations. Indeed, our functional perturbation experiments demonstrated that CG-SMG^RXFP1^ neurons control both bile and glucagon secretion but do not change intestinal food transit. CG-SMG^SHOX2^ neurons exert complementary regulations on gut transit but not secretion. This combinatorial neuron-to-organ architecture suggests that each CG-SMG population forms a functional unit to control different visceral processes. In the past decades, various models were proposed by which the sympathetic system controls peripheral organs. One such model is a uniform regulation where visceral functions are ubiquitously controlled by the sympathetic system^45,46^. Another model suggests that different organs are controlled by distinct sympathetic populations^47,48^. Our study strongly favors the latter model: specific functions are assigned to a given sympathetic population. Future studies are necessary to understand the organ- and layer-specific regulation by sympathetic populations.

### Regulation of sympathetic neuronal activity

The activity of sympathetic neurons is regulated by descending signals from the brain through pre-ganglionic neurons that reside in the spinal cord. Previous studies extensively characterized the activity of these pre-ganglionic neurons under different physiological conditions such as hunger or hypoxia^49,50^. How pre-ganglionic neurons connect to CG-SMG and other sympathetic ganglia to regulate activity is unknown. Exploring the descending sympathetic circuits at cell-type-specific precision presents an intriguing prospect, particularly after identifying genetically defined CG-SMG populations. Deciphering individual pathways may provide a means to regulate physiological body functions independently, for example, by altering blood glucose levels without affecting GI motility.

## Acknowledgments

We thank the members of the Oka laboratory, Yijie Zhang and Takako Ichiki for helpful discussion and comments; Jacob Hauser, Mari Oka and Avedis Tufenkjian for maintaining and genotyping mouse lines. TW thanks Yijie Zhang for helping with data analysis. We thank Qilan Liang and Yiping Chen for generously providing the SHOX2-Cre line; Bei Zhang and Kirsten Frieda for help on seqFISH experiments; James Linton, Felix Horns and Michael Elowitz for sharing the cell sorter; David Anderson and the Single-Cell Profiling and Engineering Center for instrumental support with scRNA-seq experiments; Wenfei Han and Ivan De Araujo for advice on surgical techniques. This work was supported by Startup funds from the President and Provost of California Institute of Technology and the Biology and Biological Engineering Division of California Institute of Technology. YO is also supported by New York Stem Cell Foundation, NIH (R01NS109997, R01NS123918), the Alfred P. Sloan Foundation, the Edward Mallinckrodt Foundation, and Heritage Medical Research Institute.

## Author contributions

TW and YO conceived the research program and designed experiments. TW performed transcriptomic, genetic, behavioral, and anatomical experiments and analyzed the data. BT performed electrophysiological characterization of CG-SMG neurons. DRY and WG designed and fabricated a microfluidic device for bile measurement. TW and YO wrote the paper. YO supervised the entire work.

## Contact Information

The authors declare no competing financial interests. Correspondence and requests for materials should be addressed to YO.

## Extended Data Figure legends

**Extended Data Fig. 1.**
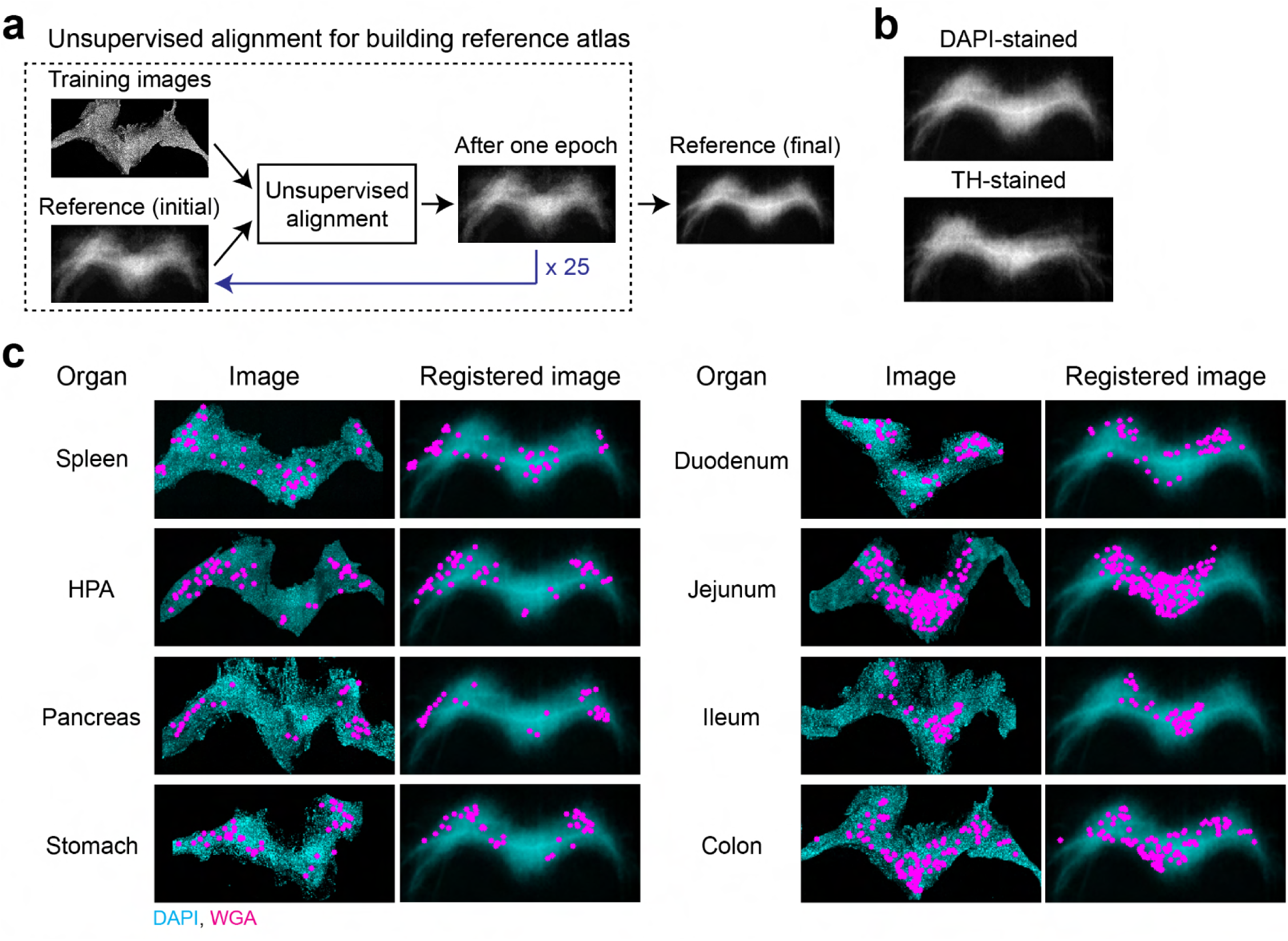
A computational pipeline for topological mapping of visceral organs. **a,** A schematic for building reference atlas through unsupervised alignment processing. **b,** Two reference images generated from DAPI-stained (top) or TH-stained (bottom) training images. **c,** Representative images exemplifying the auto-detected and registered WGA-positive cells (magenta) with DAPI staining background (cyan) for eight organ sites as indicated.

**Extended Data Fig. 2.**
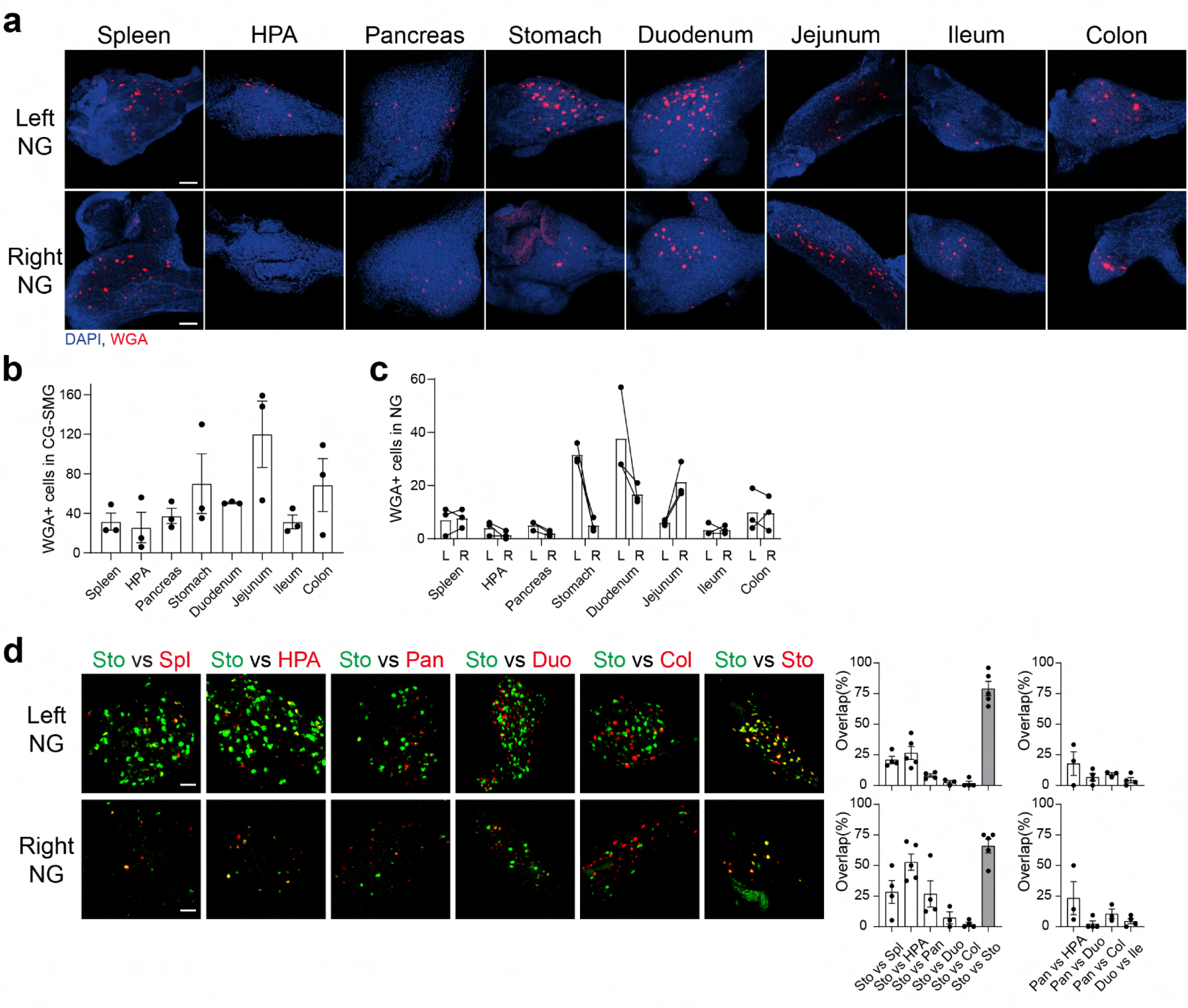
WGA retrograde tracing quantifications. **a,** Representative WGA-labeled cells (red) in the left and right nodose ganglia (NG) traced from the eight organ sites as indicated (n = 3 mice per group). **b, c,** Quantifications of WGA-positive cell number in the CG-SMG (**b**), left and right NG (**c**). **d,** Representative images and quantification of WGA-labeled cells in the left (top) and right (bottom) NG from dual-color organ tracing as indicated. n = 4, 5, 4, 3, 4, 5 mice for Sto vs Spl, Sto vs HPA, Sto vs Pan, Sto vs Duo, Sto vs Col, and Sto vs Sto. n = 3, 4, 3, 4 mice for Pan vs HPA, Pan vs Duo, Pan vs Col, and Duo vs Ile. Scale bar, 100 μm. Data are shown as mean ± s.e.m.

**Extended Data Fig. 3.**
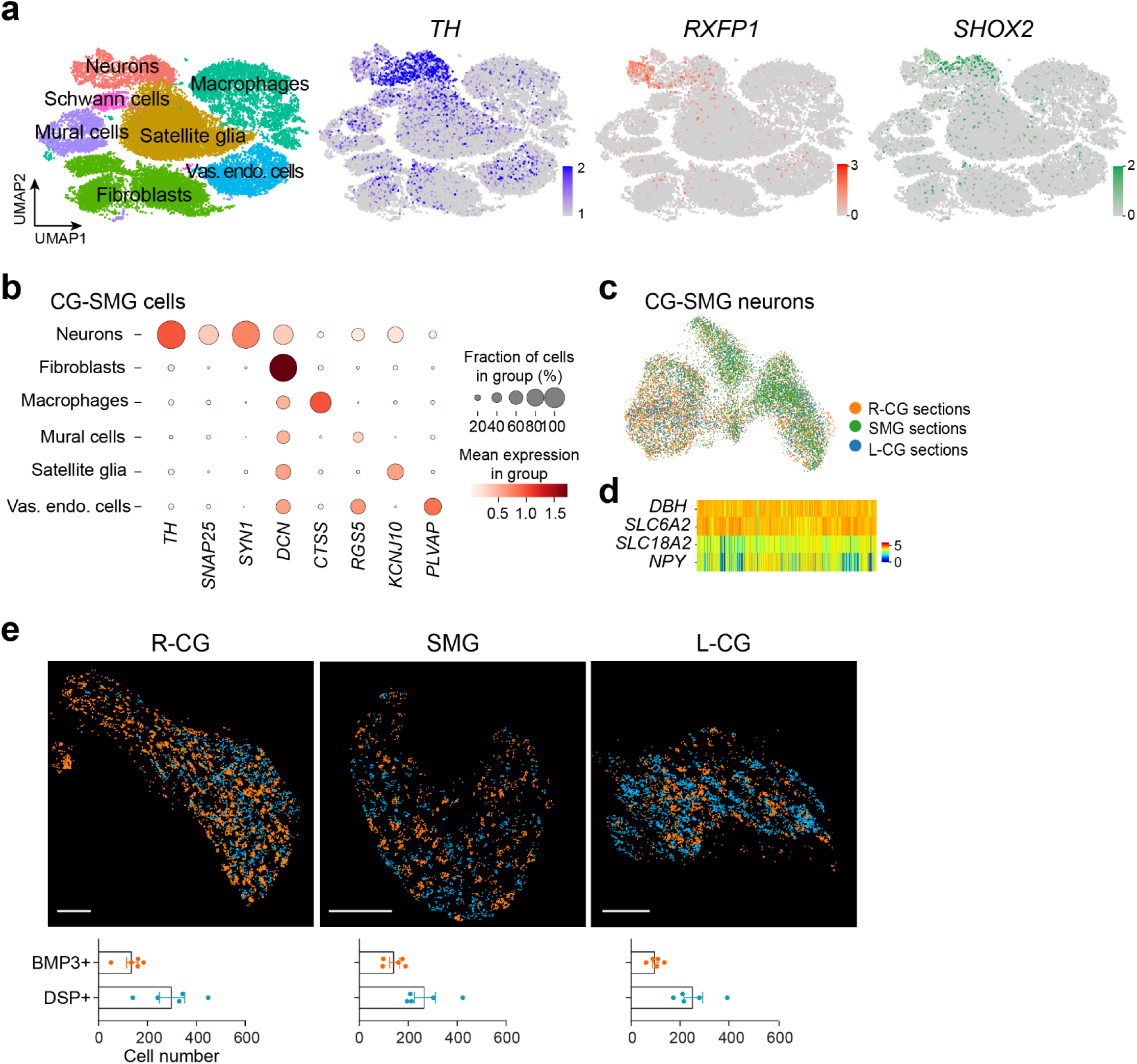
Multi-transcriptomic analyses of CG-SMG cells. **a,** ScRNA-seq analysis of CG-SMG cells. Left: UMAP embedding of CG-SMG major cell types (n = 24,873 cells). Vas. endo. cells, vascular endothelial cells. Middle and right: UMAP embedding of log-normalized *TH* (blue), *RXFP1* (red) and *SHOX2* (green) expression. **b,** Dot plot of cell-type-specific expression in CG-SMG major cell classes from seqFISH dataset. Dot size is proportional to the percentage of cells with transcript count >0 expression. Color scale represents normalized average gene expression. **c,** Heatmap of sympathetic neuronal marker gene expression in CG-SMG neurons. **d,** UMAP embedded seqFISH data from different CG-SMG areas with Harmony integration (n = 13,105 cells). **e,** Spatial localization of CG-SMG^BMP3^ and CG-SMG^DSP^ neurons. Representative seqFISH images for the expression of *BMP3* (orange) and *DSP* (cyan) in CG-SMG areas (top) and quantification of BMP3- and DSP-positive neurons (bottom, n = 5 sections of 3 mice per R-CG, SMG, L-CG). Scale bar, 200 μm.

**Extended Data Fig. 4.**
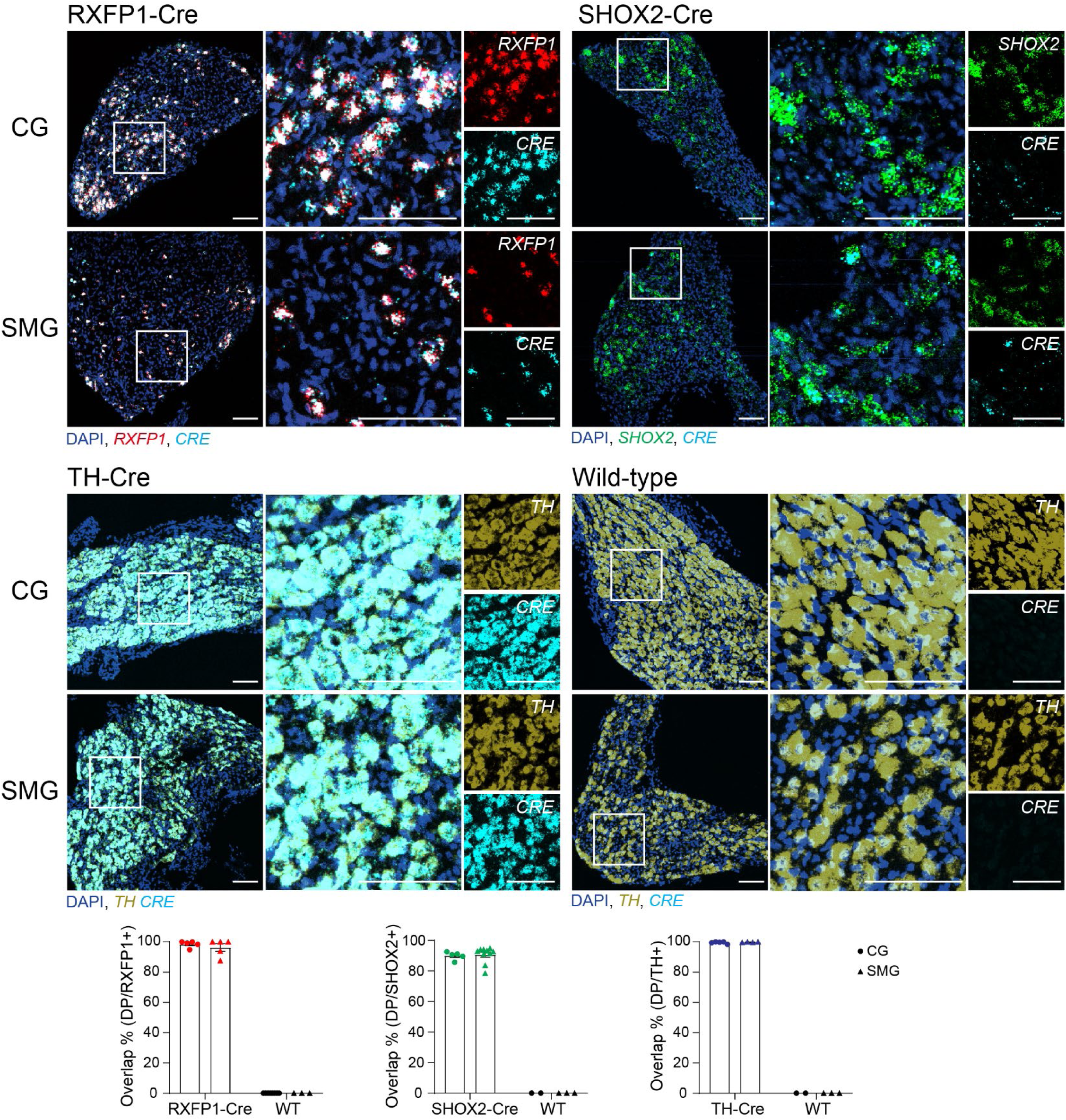
Two-color *in situ* hybridization for validating Cre expression of transgenic lines. Representative images of *Cre* and endogenous gene expression in the CG and SMG samples of RXFP1-Cre, SHOX2-Cre, TH-Cre and wild-type (WT) animals as indicated. Nuclei were visualized by DAPI staining (blue). Data were quantified from more than two mice per group. DP, double-positive. n = 5, 5, 8, 3 slices for RXFP1-Cre CG, RXFP1-Cre SMG, WT CG, and WT SMG. n = 5, 10, 2, 3 slices for SHOX2-Cre CG, SHOX2-Cre SMG, WT CG, and WT SMG. n = 5, 4, 2, 3 slices for TH-Cre CG, TH-Cre SMG, WT CG, and WT SMG. Scale bar, 100 μm. All data are shown as mean ± s.e.m.

**Extended Data Fig. 5.**
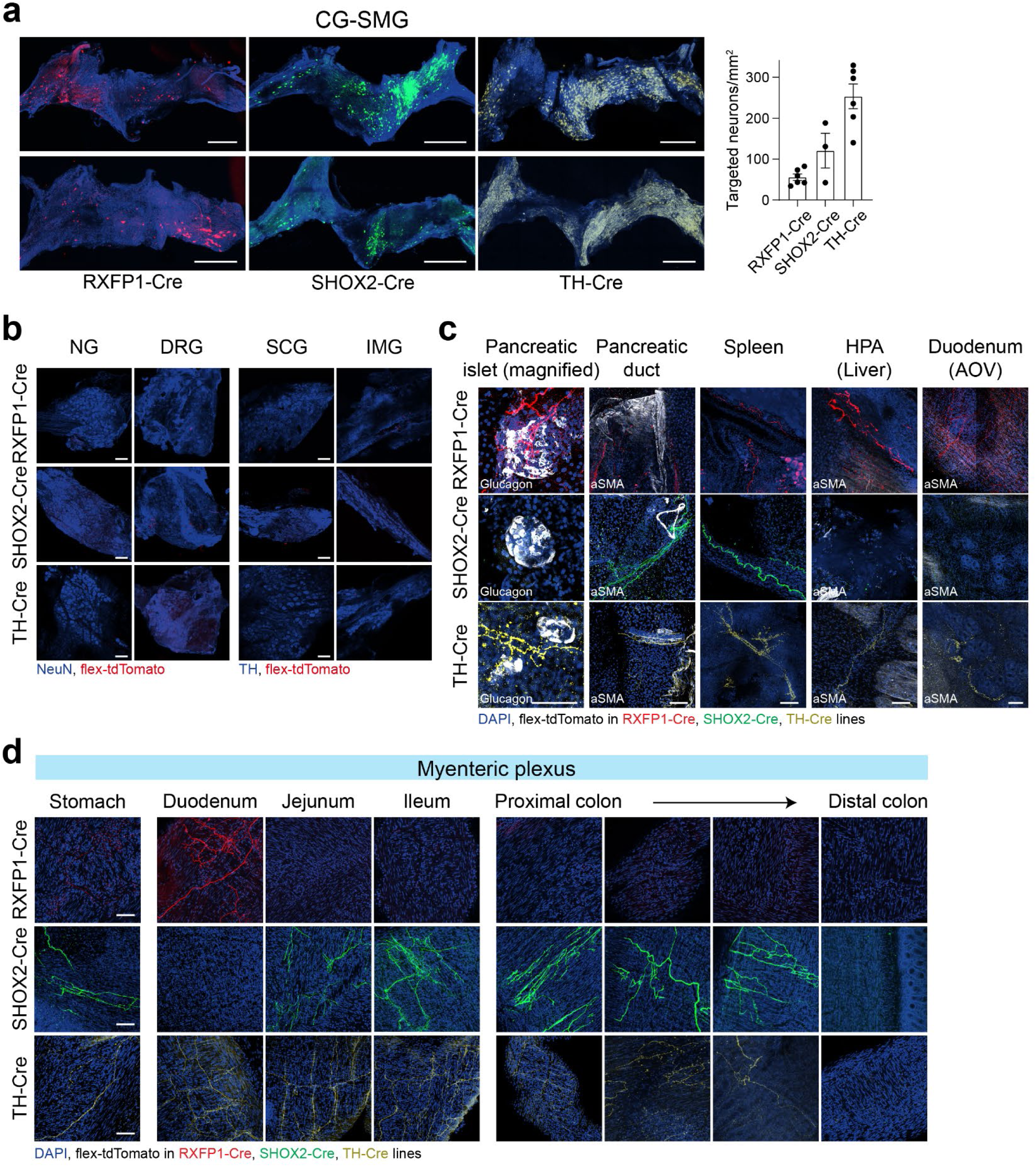
CG-SMG neural innervation of visceral organs. **a,** Representative CG-SMG images for RXFP1-, SHOX2-, and TH-targeted neurons by AAV-FLEX-tdTomato with TH staining (blue). Scale bar, 500 μm. The number of targeted neurons is quantified (right, n = 6, 3, 6 mice for RXFP1-Cre, SHOX2-Cre, TH-Cre). **b,** Peripheral ganglia histology for negative control of CG-SMG AAV injection. NG, nodose ganglion; DRG, dorsal root ganglion (T12); SCG, superior cervical ganglion; IMG, inferior mesenteric ganglion. Scale bar, 100 μm. **c,** Representative images for whole-mount organ innervation of CG-SMG^RXFP1^, CG-SMG^SHOX2^, CG-SMG^TH^ neurons. Glucagon or alpha smooth muscle actin (aSMA) staining is indicated on the images. Scale bar, 100 μm. **d,** Representative images for the innervation of the myenteric plexus on the GI tract. Nuclei were visualized by DAPI staining (blue). Scale bar, 100 μm. Data are presented as mean ± s.e.m.

**Extended Data Fig. 6.**
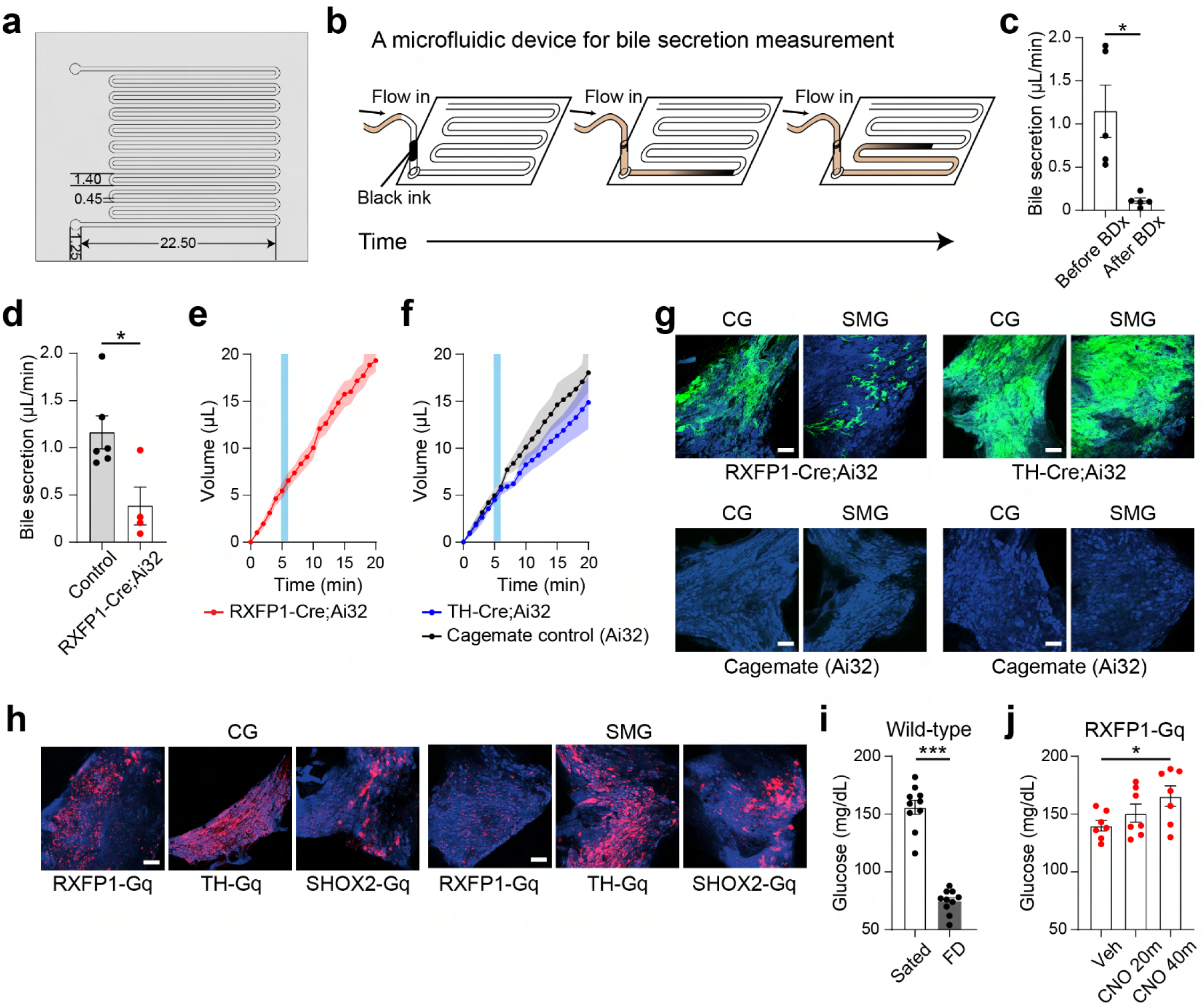
Measurements of secretory processes. **a,** A top-down view of the microfluidic device, annotated with specifications in millimeters. **b,** A diagram of bile flow in the microfluidic chamber with tip stained by the black ink for automated fluid volume detection. **c,** Average bile flow rate before and after bile duct cut (BDx, n = 5 mice). **d,** Quantified average bile flow rate during light stimulation (n = 4 for RXFP1-Cre;Ai32 mice, n = 6 for cagemate control mice). **e,** Unchanged bile production under optogenetic activation of CG-SMG^RXFP1^ neurons (n = 4 mice). Tubing was inserted to bile duct for direct measurement of bile production. **f,** Inhibition effects of stimulating CG-SMG^TH^ neurons on bile secretion (n = 3 TH-Cre;Ai32 mice, n = 2 cagemate control mice). Light pulses of 2 ms at 20 Hz were applied at CG-SMG for 1 min as indicated by the blue shade. **g, h,** Representative histological images for optogenetic (**g**) and chemogenetic (**h**) experiments as indicated. Robust ChR2-EYFP (green) or Gq-mCherry (red) expression was confirmed with sympathetic neuron marker TH staining (blue). Scale bar, 100 μm. **i,** Systemic glucose level under sated and 24-hour food-deprived (FD) states (n = 10 mice each). **j,** Effects of activating CGSMG^RXFP1^ neurons on glucose homeostasis. Systemic glucose level was measured 20 min after intraperitoneal PBS (Veh) injection, as well as 20 min (CNO 20m) or 40 min (CNO 40m) after intraperitoneal CNO administration (n = 7 mice). *P < 0.05, ***P < 0.001 by two-tailed unpaired *t*-test or one-way ANOVA with Dunnett’s multiple comparisons test. Data are presented as mean ± s.e.m.

**Extended Data Fig. 7.**
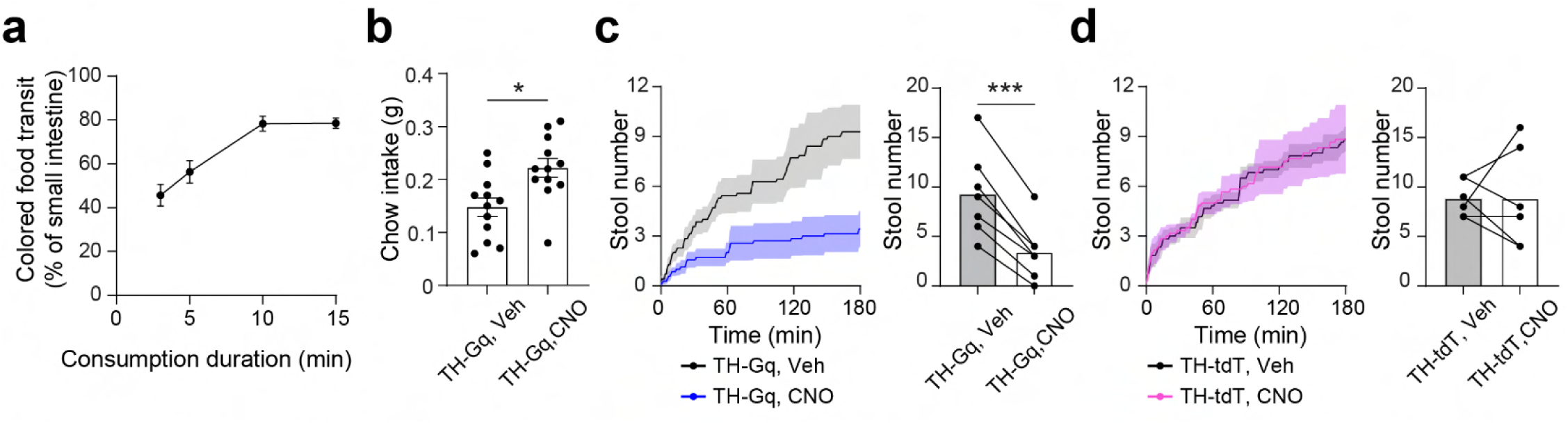
Measurements of GI transit. **a,** Quantification of colored food transit after different feeding durations as indicated (n = 5 mice per group). **b,** Chemogenetic activation of CG-SMG^TH^ neurons increased food consumption within the initial 10-min feeding after deprivation (n = 12 mice). **c,** Inhibition effects of chemogenetic stimulation of CG-SMG^TH^ neurons on stool expulsion. The number of stools after either intraperitoneal PBS (Veh) or CNO injection is plotted (left) and quantified (right, n = 7 mice). **d,** Spontaneous stool defecation of TH-Cre control animals with AAV-FLEX-tdTomato injection to CG-SMG (n = 6 mice). *P < 0.05, ***P < 0.001, or not significant by two-tailed paired *t*-test. Data are presented as mean ± s.e.m.

## Materials and Methods

### Animals

All experimental procedures were in accordance with US National Institutes of Health guidelines for the care and use of laboratory animals, and were approved by the California Institute of Technology Institutional Animal Care and Use Committee (IACUC; protocol #1694 and 1866). Male and female animals of at least 7 weeks old were used for data collection. Ai32 (#024109), and C57BL/6J (#000664) lines were acquired from the Jackson Laboratory. RXFP1-Cre line was developed in our previous publication^51^. SHOX2-Cre animals were generously provided by Y. Chen^37^. TH-Cre mice were a gift from D. Anderson. Animals were housed in a facility with controlled temperature (71-75°F) and humidity (30-70%), following a 13:11 hour light-dark cycle. Animals had *ad libitum* access to chow and water.

### Viral constructs

The following viral constructs were used: AAV9-CAG-Flex-tdTomato, 2.8 x 10^13^ genome copies per ml (Addgene, 28306-AAV9). AAV9-CAG-DIO-hM3D(Gq)-mCherry, 1.1 x 10^13^ genome copies per ml (Canadian Neurophotonics Platform Viral Vector Core Facility, 651-aav9).

### Surgery

For all survival surgeries, mice were anesthetized with a mixture of ketamine (100 mg/kg body weight) and xylazine (5 mg/kg body weight) solution via intraperitoneal administration. Ketoprofen (5 mg/kg body weight) and buprenorphine XR (3.25 mg/kg body weight) were subcutaneously applied prior to surgery. Animals were placed on their backs and the abdomen was incised along the midline.

### Celiac-superior mesenteric ganglia injection

The CG-SMG complex was exposed following reflection of the viscera. The small and large intestine were externalized and covered by a wet, sterile gauze. Using the superior mesenteric artery and celiac artery as landmarks, 100-300 nl virus containing 0.05% Fast Green FCF (Sigma, F7252-5G) was delivered to three sites (left CG, SMG and right CG) of the CG-SMG complex at a rate of 100 nl/min with a microprocessor-controlled injection system (World Precision Instruments, Nanolitre 2000). Successful injection was visualized by Fast Green FCF dye slowly filling the ganglion without leakage. For the anterograde virus tracing experiments, tissue collection was done at least 3 weeks after the injections. For the chemogenetic experiments, animals had at least 2 weeks of recovery period.

### Retrograde tracing from visceral organs

After the abdominal incision, a total of 0.5-1 μl retrograde tracer (WGA 488 or WGA 647, ThermoFisher, W11261, 5 mg/mL) was injected to the target organ at 100 nl/min. For GI tract injection, tracer was delivered into the layer between the muscularis externa and serosa at 4-8 sites, covering a total area of approximately 1 cm^2^. For injection to the spleen and pancreas, tracer was uniformly administered across the organ’s surface at 4-8 evenly spaced sites. For injection to the HPA, tracer was delivered to the connective tissue surrounding the portal vein and the areas of the liver where portal vein enters. Peripheral ganglia were harvested three days post-injections for histological and imaging analyses. For dual-color tracing experiments, two or three regions containing both WGA 488- and WGA 647-labeled cells were quantified for the ratio of double-positive cells among all WGA-positive cells.

### Organ topological mapping

To perform organ topological mapping on the CG-SMG complex, CG-SMG neurons were retrogradely traced from eight organ sites (spleen, HPA, pancreas, stomach, duodenum, jejunum, ileum and colon) by WGA dye as described above. Three mice per organ site were injected. We developed a computational pipeline to build a standardized reference anatomy image, align WGA-labeled ganglia image, and register WGA-positive cells, by applying the VoxelMorph library (version 0.2)^52,53^. In brief, the entire CG-SMG ganglia complex was whole-mount and z-stack imaged to visualize all projecting neurons via confocal microscopy (Leica, TCS SP8). By digitally drawing vertical lines from both the left and right sides towards the center, we determine the lateral boundaries to be at 20% of its maximum width.

The atlas was generated using the TemplateCreation network with default architecture and mean squared error (MSE) loss. Other model settings were: batch size = 1, steps per epoch = 100, loss weight = [0.5, 0.5, 1, 0.01], learning rate = 1e-4. All input images for the network were first resized to 128 pixels x 64 pixels. In particular, the initialization of the reference was the average of the training dataset comprising twenty stained CG-SMG training images. We trained the neural network over 25 epochs for performance optimization. Consistent atlas was built from DAPI- or TH-stained images (Extended Data Fig. 1b).

The atlas generated from the CG-SMG dataset of DAPI staining was used for the following image registration on the organ mapping dataset of 24 WGA-labeled images. WGA-positive cells on DAPI background were auto-detected using the OpenCV library, and the cell coordinates were obtained. The number of WGA-positive cells was automatically counted (Extended Data Fig. 2b). Next, we trained the VxmDense network on WGA-labeled images with default architecture, MSE loss, batch size = 4, steps per epoch = 50, loss weight = [1, 0.1], learning rate = 1e-4, and number of epochs = 25. All input images for the network are first resized to 256 pixels x 128 pixels. Each WGA-labeled image was aligned to the reference atlas based on DAPI staining by the unsupervised model. This same spatial transformation was applied to WGA-positive cells. We calculated the mapping of WGA-positive cell coordinates from WGA-labeled images to the atlas. After image registration, WGA-positive cells from three images were plotted on the atlas image as scaled Gaussian heatmaps.

### Bile secretion measurement

#### Fabrication of the microfluidic chamber

The microfluidic channels were 3D printed (Elegoo Mars 3) using ultraviolet-sensitive polymer resin (Anycubic High Clear Resin). The structure was then cleaned with isopropyl alcohol to remove excess resin, followed by ultraviolet treatment under 365 nm light for 2 min and heat curing at 80°C for 2 min. The chamber’s top layer, a 75 µm polyethylene terephthalate (PET, McMaster-Carr) sheet, was attached using double-sided medical adhesive (3M). This bilayer was cut using a 50W CO2 laser cutter (Universal Laser System) at 40% power, 100% speed, 1000 PPI, vector mode, 2 copies. The PET layer, with laser-cut inlet and outlet holes, was then laminated onto the 3D printed channels. A watertight inlet seal was created by gluing a silicone rubber connector (Dragon Skin^TM^ FX-Pro^TM^, Smooth-On), which was shaped in a 3D printed mold and cured at 80°C for 5 min.

#### Surgical procedure for bile secretion measurement

Under isoflurane anesthesia, a skin incision was made along the abdominal midline. A small fragment (< 1 cm, within 1.5 cm from the sphincter) of duodenum containing the bile duct entry was isolated by tying both ends with sutures (Ethicon, K802H). Then an intraintestinal tube (HelixMark®, 60-011-04) was positioned adjacent to the bile duct entry site, secured by sutures, and connected to the customized microfluidic chamber. For the bile duct tubing, the duct was first ligated near the duodenum and then incised for the tube (Instech, BTPU-014) insertion. A camera was positioned vertically above the microfluidic chamber for video recording at a frame rate of 30 Hz.

For optogenetic stimulation, an optic fiber was held with tip close to the right CG, delivering 20 Hz, 473-nm laser pulses (2-ms duration) for 1 min with the pulse generator (Quantum composers, Sapphire 9200). The laser intensity was maintained at 5 mW at the fiber tip. For chemogenetic manipulation, clozapine N-oxide (CNO) (Sigma, 2.5 mg/kg body weight) or vehicle (PBS) was locally administered around the hepatic region (bile duct and duodenum). Mice were immediately euthanized for histological verification after the recording session.

#### Computer vision-based video analysis

Dried soluble black ink was placed at the inlet of the microfluidic chamber to visualize the fluid flow’s tip (Extended Data Fig. 6b). As bile enters the chamber, the tip of the flow becomes marked by the black ink, enhancing its visibility. The initial 10-min equilibration period was not included in the video analysis. The fluid flow tip’s location and the cumulative area covered by the fluid were tracked using background subtraction method in OpenCV. The volume of fluid over time was calculated based on the specifications of the microfluidic device, and this data was recorded for every frame, achieving a resolution at the nanoliter scale with a sampling rate of 30 Hz. To normalize the flow volume in Fig. 4e and f, the average flow rate in Fig. 4d was subtracted. Data points representing the first frame of each minute were plotted unless indicated in the figure legend. For optogenetic experiments, the average flow rates during the minute preceding light exposure and during the minute of light stimulation were quantified. For chemogenetic activation experiments, the average flow rates during the 5 min before and after CNO application were quantified.

### Plasma glucagon concentration measurements

The following groups were tested: sated, 24-hour food-deprived wild-type mice, and sated transgenic mice with DREADD-expressing CG-SMG neurons. Intraperitoneal CNO injection was performed 20 min before blood collection. For food deprivation, animals had free access to water and single chow pellet daily. The hepatic portal vein blood was collected to tubes with sodium fluoride / Na_2_EDTA additive (BD, 365992) under isoflurane anesthesia. Plasma was obtained by centrifugation at 1500 g for 20 min. The glucagon concentration was determined by ELISA kit (RayBiotech, EIAM-GLU-1).

### Systemic glucose level measurements

Systemic glucose level was assessed using a glucometer (KTO-MOJO) on blood samples obtained following toenail trimming under isoflurane anesthesia. Sated and 24-hour food-deprived wild-type mice were examined. RXFP1-Gq animals were tested 20 min after intraperitoneal injection of either PBS (as a control vehicle) or CNO. Additional measurements were conducted 40 min post-CNO delivery. Each measurement was separated by a minimum recovery period of one week.

### Assessment of food transit

Animals were food deprived for 24 hours prior to the food transit experiments. Chow pellets were uniformly coated with food-grade dye through a quick immersion process, followed by drying on a hotplate at 80 °C for 20 min. Twenty mins following the intraperitoneal administration of PBS or CNO (2.5 mg/kg body weight), animals were placed in an empty cage with one colored pellet and were euthanized between 3 to 15 minutes into their meal, as indicated in figures. The gut was immediately evaluated either by terminal dissection or through rapid surgical opening under isoflurane anesthesia. The extent to which the dye had traveled was quantified as a percentage of the total length of the small intestine, from the pyloric sphincter to the ileocecal junction.

### Behavioural assays

#### Stool analysis

Individual sated animals were located in a 25 cm x 25 cm acrylic open field chamber after intraperitoneal PBS or CNO injection, and were recorded for three hours at 30 frames per sec. The stool number was quantified based on the videos.

#### Food intake

Animals were individually housed and acclimated to the BioDAQ cages (Research Diets) for 3 consecutive days. For food intake measurement, mice were food deprived for 24 hours and intraperitoneally injected PBS or CNO 20 min before the experiments. The total chow intake in 10 min was quantified.

### CG-SMG neuron electrophysiological recordings

#### CG-SMG neuron isolation

CG-SMG neurons from adult mice were acutely isolated using an enzymatic dispersion technique modified from a previous procedure^54^. In brief, mice were euthanized by cervical dislocation under isoflurane anesthesia, and perfused with cold artificial cerebrospinal fluid (ACSF). The CG-SMG complex was dissected to cold ACSF, and the connective tissue capsule was carefully peeled and removed under dissecting microscope. The ganglia were minced with iridectomy scissors into fine pieces and then transferred to a 35 mm petri dish containing 2 ml enzyme mix (0.4 mg/ml Trypsin (Roche), 1.2 mg/ml collagenase D (Roche), 0.15 mg/ml DNase I (Roche) dissolved in EBSS (Sigma)). The tissue fragments were incubated in the enzyme mix for 1 hour at 35 °C, bubbled with 5% CO2 and 95% O2. After enzyme digestion, 2 ml of minimum essential medium (MEM) containing 10% FBS, 1% glutamine and 1% penicillin-streptomycin antibiotics was added to the petri dish. The cell suspension was then centrifuged at 200 g for 5 min. Discard the supernatant and resuspend the cells with the MEM solution mentioned above. Gentle trituration is optional to help break down ganglion tissue and isolate CG-SMG neurons. The cells were plated onto cover glass coated with Cell-tek (Corning) and incubated in a humidified atmosphere at 5% CO2 in air at 37 °C prior to patch-clamp recordings.

#### CG-SMG neuron electrophysiology

For patch-clamp recordings, isolated CG-SMG neurons were plated in recording chamber and placed on an upright microscope (Examiner.D1, Zeiss) perfused with normal ACSF (in mM: NaCl 124, KCl 2.5, NaH_2_PO_4_ 1.2, NaHCO_3_ 24, glucose 25, MgSO_4_ 1, CaCl_2_ 2, and bubbled with 95% O_2_/5%CO_2_). Electrical signals were filtered at 3kHz with Axon MultiClamp 700B (Molecular Devices) and collected at 20 kHz with Axon Digidata 1550A (Molecular Devices). For current clamp recordings, intracellular solution contains (in mM) K-gluconate 145, NaCl 2, KCl 4, HEPES 10, EGTA 0.2, Mg-ATP 4, Na-GTP 0.3 (pH 7.3). Thin wall patch pipettes (OD = 1.5 mm, ID = 1.17 mm, Sutter Instrument) were fabricated on a model P-97 micropipette puller (Shutter Instrument) and fire polished on a microforge to a resistance of 2 – 3 MΩ.

For optogenetic experiments, light beam from an LED light source (X-Cite 120LED, Excelitas Technologies) was delivered through an optical filter (475/30). Light pulses (10 ms) were given at 2 – 20 Hz for 4 sec in experiments as in Fig. 4b to evoke action potentials in CG-SMG neurons. For chemogenetic experiments, CNO was applied through a puffing pipette directly located adjacent to the recording neuron.

### Single-cell RNA sequencing

Five male and five female C57BL/6J mice of 7 weeks old were used to prepare per single-cell RNA sequencing (scRNA-seq) run on the 10x Genomics platform. Across two sequencing runs, a total of 24,873 cells were profiled, which included 1,558 neurons. Upon isoflurane anesthesia, mice were euthanized by cervical dislocation, and perfused with ice-cold HEPES-ACSF (in mM: NaCl 92, KCl 2.5, NaH_2_PO_4_ 1.25, NaHCO_3_ 30, HEPES 20, glucose 25, MgSO_4_ 1, CaCl_2_ 2, kynurenic acid Na salt 1, Na-ascorbate 5, thiourea 2, Na-pyruvate 3, and bubbled with 95% O_2_/5%CO_2_). Under dissecting microscope, the CG-SMG complex was rapidly extracted to 1ml ice-cold ACSF, and the connective tissue capsule was carefully peeled and removed. After tissue harvest, HEPES-ACSF was replaced by 1 ml enzyme digestion buffer 1 (HEPES-ACSF) containing papain (80 U/ml; Worthington, PAPL, LS003119; pre-activated with 2.5 mM cysteine and a 30-min incubation at 34 °C) and TrypLE™ Express Enzyme (1X, 400 μl/ml; ThermoFisher, 12604013). During enzymatic digestion the tissue was pipetted periodically every 10 min. After incubation at 34 °C with gentle carbonation for 35 min, the CG-SMG tissue was carefully washed one time with 1 ml HEPES-ACSF. One ml of the enzyme digestion buffer 2 (HEPES-ACSF) containing collagenase type 2 (2 mg/ml; Worthington, LS004174), dispase II (2 mg/ml; Sigma, D4693-1G), deoxyribonuclease I (0.2 mg/ml; Worthington, LS002007) and supplemented with 2 mM CaCl2 was applied. After incubation at 34 °C with gentle carbonation for 20 min, the CG-SMG tissue was carefully washed one time with 1 ml HEPES-ACSF. The medium was replaced with 100 μl HEPES-ACSF containing 0.1 μM Calcein AM (ThermoFisher, C1430), 4 μM ethidium homodimer-1 (EthD1, ThermoFisher, E1169), and 0.02 mg/ml deoxyribonuclease I. Tissue was gently triturated into a single-cell suspension with consecutive rounds of trituration with fire-polished glass Pasteur pipettes with tip diameters of around 600, 300 and 150 μm. The suspension was brought up to 200 μl and pipetted through a 40-μm cell strainer (pluriStrainer, 43-10040) into a new microcentrifuge tube and incubated on ice for 5 min. Calcein AM-positive and EthD1-negative singlets were sorted (Sony, MA900) and collected in 4 °C HEPES-ACSF containing 0.05% BSA. The sorted cell suspension was centrifuged with 400 g for 4 min at 4 °C. The supernatant was discarded, and the cell pellet was resuspended with 55 μl fresh ice-cold HEPES-ACSF and kept on ice while the cell density was counted with a hemocytometer. The final cell suspension volume estimated to retrieve 10,000 single-cell transcriptomes was loaded to the 10X Single Cell G chip (10x Genomics, PN-1000127). The Chromium Single Cell 3’ GEM, Library and Gel Bead Kit v3.1 (PN-1000128) and the Single Index Kit T Set A (PN-1000213) were used. The cDNA and library amplification underwent 11 and 12 cycles respectively. The scRNA-seq libraries were sequenced on a NovaSeq S4 lane (paired-end 150). The sequencing reads were mapped to the custom pre-mRNA reference transcriptome^55^, and gene-cell matrices were generated via the 10x Genomics Cell Ranger v.6.1.2 pipeline. Subsequent gene expression analyses were conducted in Python (3.9.13) using ScanPy (1.9.5) as previously described^56^. Cells were filtered if possessing fewer than 1,500 or more than 45,000 unique transcripts, or more than 10% of mitochondrial transcripts.

### Spatial transcriptomics

#### SeqFISH gene panel design

A custom seqFISH panel was designed to incorporate CG-SMG cell-type-marker genes based on our scRNA-seq data analysis. The panel contained 371 genes of which 341 were detected using barcoded seqFISH imaging and 30 were identified serially via single-molecule FISH. The panel was custom ordered by Spatial Genomics, Inc.

#### SeqFISH sample preparation

Adult (8-12 weeks old) C57BL/6J male mice were euthanized by cervical dislocation under isoflurane anesthesia. After dissecting the CG-SMG complex to ice-cold PBS, right, left and medial CG-SMG samples were obtained by surgically separating the connected CG-SMG region between the celiac artery and the superior mesenteric artery. The tissue was freshly embedded in Tissue-Tek O.C.T. Compound (Sakura, 4583), and then flash-frozen in liquid nitrogen. Cryosections of 10-μm thickness were cut and mounted onto treated coverslips. Immediately post-collection, sections were fixed in fresh 4% paraformaldehyde (PFA, Thermo Scientific, 28908) for 15 min at room temperature. After rinsing three times in 1X PBS for 5 min each, sections were dehydrated using 70% ethanol for 30 sec at room temperature. The sections were air-dried at room temperature for approximately 15 min and stored at −80 °C. SeqFISH experiments were performed using the seqFISH+ protocol with some modifications at Spatial Genomics, Inc. Briefly, the fixed tissue sections underwent permeabilization in 70% ethanol followed by clearing, rinsing, and air-drying steps. The coverslip containing each section was assembled into Spatial Genomics custom flow cells. The flow cells were then hybridized with the seqFISH primary probe panel and incubated in a humidified chamber at 37 °C for 24 hours. After hybridization, samples were washed with buffers for subsequent imaging.

#### SeqFISH imaging via Gene Positioning System

Imaging was performed using the Gene Positioning System (GenePS, Spatial Genomics, Inc.), which enables automated image acquisition, reagent delivery, and data processing. The ganglion section was selected as a region of interest (ROI) for the experiment based on the brightfield and/or DAPI images. Automated experiment execution was initiated post-ROI selection. Each experiment proceeded through multiple rounds of decode probe hybridization, imaging, and signal removal until all the hybridization rounds were complete.

#### Image processing and analyses

Raw-image files were processed on instrument to align images across multiple hybridization rounds and detect RNA fluorescent signals. The data were further analyzed using custom Spatial Genomics analysis software to decode the transcript identities and segment cells. Cell segmentation was performed using a machine learning algorithm based on the cell nuclear DAPI stain or TH stain. The decoded RNA molecules were then assigned to individual cells, generating cell-by-gene count matrices and individual cell center coordinates for each ROI.

Subsequent gene expression analyses were conducted in Python (3.9.13) using ScanPy (1.9.5) as previously described. Cells that had unique gene counts of less than 10 were filtered. Harmony^57^ package was used to integrate different batches of seqFISH datasets. Gene expression count data were normalized per cell and log-transformed for downstream analyses by sc.pp.normalize_total and sc.pp.log1p functions. We then performed dimensionality reduction with principal component analysis (PCA) and uniform manifold approximation and projection (UMAP) analysis, followed by unsupervised clustering using the Leiden algorithm through the sc.tl.leiden function (resolution parameter set to 1). To identify major cell classes, we used a DAPI-based segmented dataset and compared the expression patterns of known cell-type specific marker genes within each cluster. To reduce the effects of high *TH* expression in non-neuronal cells on clustering results, we prioritized the annotation of non-neuronal cells over neurons and accordingly adjusted the weighing of *TH* expression level in non-neuronal cells. To determine neuron subtypes, we proceeded with 13,105 neurons from TH-based segmentation. The neuronal subtypes (resolution parameter ranging from 0.1 to 0.5) were hierarchically clustered based on Pearson correlation using the sc.tl.dendrogram function. To quantify the number of neurons, RXFP1-, SHOX2-, TH-, BMP3-, and DSP-positive neurons are defined as log-normalized expression RXFP1 > 0.6, SHOX2 > 0.6, TH > 0, BMP3 > 0, DSP > 0, respectively.

### RNAscope-based multicolor *in situ* hybridization

*In situ* hybridization was performed with the RNAscope Multiplex Fluorescent Assay version 2 (ACD, 323110). Fresh-frozen lateral and medial CG-SMG ganglia sections from C57BL/6J, RXFP1-Cre, SHOX-Cre and TH-Cre animals were prepared following the manufacturer’s protocols. The following probes were applied to visualize gene expression: CRE (ACD, 312281, 474001), RXFP1 (ACD, 458001, 458001-C3), SHOX2 (ACD, 554291-C3), and TH (ACD, 317621-C2). The samples were imaged with confocal microscopy and were manually quantified.

### Histology

#### Ganglia expression

Mice were euthanized with CO2 and perfused with PBS followed by 4% PFA. Ganglia were extracted and post-fixed for 1 hour at 4 °C in PFA, and rinsed two times with PBS. The ganglion sample was blocked (10% donkey serum, 0.2% Triton X-100 in PBS) for 1 hour at room temperature and incubated with primary antibodies overnight at 4°C. The following primary antibodies were used: chicken anti-NeuN antibody (1:500; Millipore, ABN91), and chicken anti-TH antibody (1:500; AvesLab, TYH-0020). After washing 3 x 10 min with PBS, the tissue was stained with the secondary antibody (1:500; Jackson Immunoresearch) and DAPI (2 mg/ml) for 4 hours under room temperature. The following secondary antibodies were used: donkey anti-chicken 488 (703-545-155), and donkey anti-chicken 647 (703-605-155). After three times washing with PBS, the ganglia were mounted on the glass slide and imaged with confocal microscopy.

#### Organ innervation

Mice were euthanized with CO2 and perfused with PBS followed by 4% PFA. Tissue samples were extracted and post-fixed overnight at 4 °C in PFA. Approximately 2 cm-length samples from the middle of each segment of the GI tract, such as the duodenum, jejunum, and ileum, were harvested. The entire colon was uniformly divided into four sections and collected, from the proximal to the distal end. The tissue was then longitudinally incised and flattened. For whole-mount visceral organ staining, tissue was blocked (10% donkey serum, 0.2% Triton X-100 in PBS) overnight at 4 °C and incubated with primary antibodies overnight at 4°C. The following primary antibodies were used: goat anti-alpha-smooth muscle actin (1:1000; Novus Biologicals, NB300-978), rabbit anti-glucagon antibody (1:500; Abcam, ab92517), rat anti-mCherry (1:500; Invitrogen, M11217). After washing 5 x 1 hour with 0.1% PBST, the tissue was stained with the secondary antibody (1:500; Jackson Immunoresearch) and DAPI (2 mg/ml) overnight at room temperature, washed 3 x 1 hour with 0.1% PBST. The following secondary antibodies were used: donkey anti-rat Cy3 (712-165-150), donkey anti-goat 647 (705-605-147), and donkey anti-rabbit 488 (711-545-152). Tissue was then cleared in ScaleS solution overnight at room temperature. Samples were mounted in ScaleS with 0.5-mm spacers (SUNJin Lab, IS011) on slides and imaged with confocal microscopy.

#### Quantification of ganglia expression and nerve density

One CG and one SMG z-stack image of 775 μm x 775 μm were quantified for each animal in ImageJ. The average number of targeted cells from these two images was calculated and plotted as a unit of cell number per square millimeter in Extended Data Fig. 5a.

Projected organ innervation images of 581 μm x 581 μm were automatically segmented by the MaxEntropy auto-threshold function in ImageJ. For each organ from each animal, two images were quantified. Nerve density was calculated as the ratio of Nerve Area to Total Area as described in a recent publication^44^.

### Statistics and data collection

Data were processed and analyzed by Prism 10.2.0. No statistical methods were used to predetermine the sample sizes. The experiments were not randomized, and investigators were not blinded to allocation during experiments and outcome assessment. Data are presented as means ± s.e.m. Two-tailed paired or unpaired *t*-test, and one- or two-way ANOVA with post-hoc tests were applied to decide the significance level. No significance (ns) was set at p > 0.05. The significant threshold was held at \**P* < 0.05, \*\**P* < 0.01, \*\*\**P* < 0.001. The sample sizes and statistically significant effects are reported in each figure or figure legend.

## Data availability

Additional data that support the finding of this study are available from the corresponding author upon reasonable request.

## Code availability

All Python code used in this manuscript is available from the corresponding author upon reasonable request. For the organ topological mapping, we imported the VoxelMorph library at https://github.com/voxelmorph/voxelmorph.

